# De novo drug designing coupled with brute force screening and structure guided lead optimization gives highly specific inhibitor of METTL3: a potential cure for Acute Myeloid Leukaemia

**DOI:** 10.1101/2023.09.11.556977

**Authors:** Manisha Ganguly, Radhika Gupta, Amlan Roychowdhury, Ditipriya Hazra

**Author notes:** Correspondence Ditipriya Hazra, Amlan Roychowdhury. These authors contributed equally: Manisha Ganguly and Radhika Gupta.

## Abstract

Expression of METTL3, a SAM dependent methyltransferase, which deposits m6A on mRNA is linked to poor prognosis in Acute Myeloid Leukaemia and other type of cancers. Down regulation of this epitranscriptomic regulator has been found to inhibit cancer progression. Silencing the methyltransferase activity of METTL3 is a lucrative strategy to design anticancer drugs. In this study 3600 commercially available molecules were screened against METTL3 using brute force screening approach. However, none of these compounds take advantage of the unique Y-shaped binding cavity of the protein, raising the need for de novo drug designing strategies. As such, 125 branched, Y-shaped molecules were designed by “stitching” together the chemical fragments of the best inhibitors that interact strongly with the METTL3 binding pocket. This results in molecules that have the three-dimensional structure and functional groups which enable it to fit in the METTL3 cavity like fingers in a glove, having unprecedented selectivity and binding affinities. The designed compounds were further refined based on Lipinski’s rule, docking score and synthetic accessibility. The molecules faring well in these criteria were simulated for 100ns to check the stability of the protein inhibitor complex followed by binding free energy calculation.

## INTRODUCTION

Eukaryotic RNAs are known to be modified post-transcriptionally, adding another layer of regulation in the flow of genetic message. N6-methyladenosine more commonly known as m6A, is the most prevalent modification among the different post-transcriptional modifications found in the eukaryotic RNA, including N1-methyladenosine (m1A), methoxycarbonylmethyl-2′-O-methyluridine (mcm5Um), 5-methylcytosine (m5C) and others (X. Wang & He, 2014). Single-nucleotide-resolution mapping of transcriptome revealed that thousands of RNAs carry m6A modification (Ke et al., 2015; Linder et al., 2015; Yue et al., 2015). These modifications are enriched in the 3′ UTR near the stop codon, however, some m6A marks are also found in the 5′ UTR as well as coding regions. These studies further revealed that METTL3, a S-adenosylmethionine (SAM) dependent methyl transferase (MTase) deposits the m6A mark in the following consensus sequence, DRACH (D = A/G/U, R = A/G, and H = A/C/U) of human mRNA.

The m6A machinery is composed of several proteins: other than the SAM dependent methyltransferases or writers there are erasers or demethylases and readers or m6A binding proteins. Erasers are responsible for demethylation of the adenosine. Fat mass and obesity-associated protein (FTO) is an eraser situated both in nucleus and cytoplasm, from where it can regulate RNA modification, transcription and metabolism (Jia et al., 2011). Another example of eraser protein is Alkylated DNA repair protein ALKB homologue 5 (ALKBH5). It performs m6A demethylation and regulates RNA processing (Zheng et al., 2013). In contrast the YTH domain reader proteins, (YTHDC1, YTHDC2, YTHDF1, YTHDF2, YTHDF3), are involved in mRNA stability, translation efficiency, splicing and RNA export (Hazra et al., 2019).

Writer proteins catalyse the formation of m6A. The main writer protein in human beings METTL3 is part of a multicomponent m6A methyltransferase complex (MTC), composed of METTL3-METTL14 heterodimer core, Vir-like m6A methyltransferase-associated protein (VIRMA), and RNA binding motif protein 15 (RBM15/15B), Wilms tumour 1 associated protein (WTAP), ZC3H13 and so on (Zeng et al., 2020). The crystal structures of METTL3 and METTL14 reveal that they form a heterodimeric complex having butterfly shape in an asymmetric unit (Śledź & Jinek, 2016; P. Wang et al., 2016; X. Wang et al., 2016), where METTL14 helps in stabilizing the complex.

Recent studies showed that METTL3 serves as an oncogene which is upregulated in different types of cancers, including Acute Myeloid Leukaemia (AML) (Vu et al., 2017), Glioblastoma (J. Shi et al., 2021), lung cancer (Ma et al., 2022), Colorectal cancer (Chen et al., 2022), Prostate cancer (Yuan et al., 2020), Gastric cancer (Yang et al., 2020), Breast cancer (Y. Shi et al., 2020).

In Adult Acute Myeloid Leukaemia (AML), METTL3 plays an important role. Certain oncogenes such as PTEN, c-MYC, and BCL2 were found to be upregulated in an m6A-dependent manner and ablation of METTL3 in AML cell line reduced cancer progression and induced apoptosis (Vu et al., 2017). In colorectal cancer (CRC), METTL3 is over expressed and induces the progression of CRC metastatic tissues (Li et al., 2019b). In lung cancer it has been reported that cell progression is affected by raised level of METTL3 which induces drug resistance and progression of non-small-cell lung cancer (NSCLC) cells by upregulating the levels of m6 A modifications (Jin et al., 2023). The deletion or suppression of METTL3 expression has been found to affect tumour progression and migration of cancerous cells. Hence targeting METTL3 and designing structure specific inhibitors to silence the MTase activity could be an effective way to develop new drugs to combat with different cancers. The targeted methyl transferase domain of METTL3 is composed of 8 β strands and 7 α helices connected by loops (Figure 1A). Most of the SAM or SAH (S-adenosyl homocysteine, a non-hydrolysable analogue of SAM) binding pocket amino acid residues, ASP377, ASP395, ASN539, GLU532, ARG536, ASN549, & GLN550 are residing on loop regions where only HIS538, ASN539 are residing on a small helix, α_6_. Figure 1B represents the surface structure of METTL3 with a clear view of the Y shaped ligand binding pocket as described by the authors in their previous work (Manna et al., 2023). The binding pocket has been described as a composite structure containing a central hub region (red) connected with three sub-pockets annotated as P1 (blue), P2 (green) and P3 (yellow). The figure also describes the binding pose of SAH (dark yellow) and amentoflavone (AMF) (light grey) in the Y-shaped pocket. It clearly shows the relative occupancies of these two ligands are different in the Y shaped cavity. SAH has occupied P2 and P3 (SAM also shares the same binding pose as SAH, PDB ID 5IL1) where AMF binds with P1 and P2. This observation triggered the hypothesis that the inhibitor binding pose for METTL3 is partially different from the co-factor binding site and may provide unique insight in designing target specific inhibitor. The only commercially available inhibitor of METTL3, STM2457, also occupies P1 and P2 pocket.

**Figure 1:**
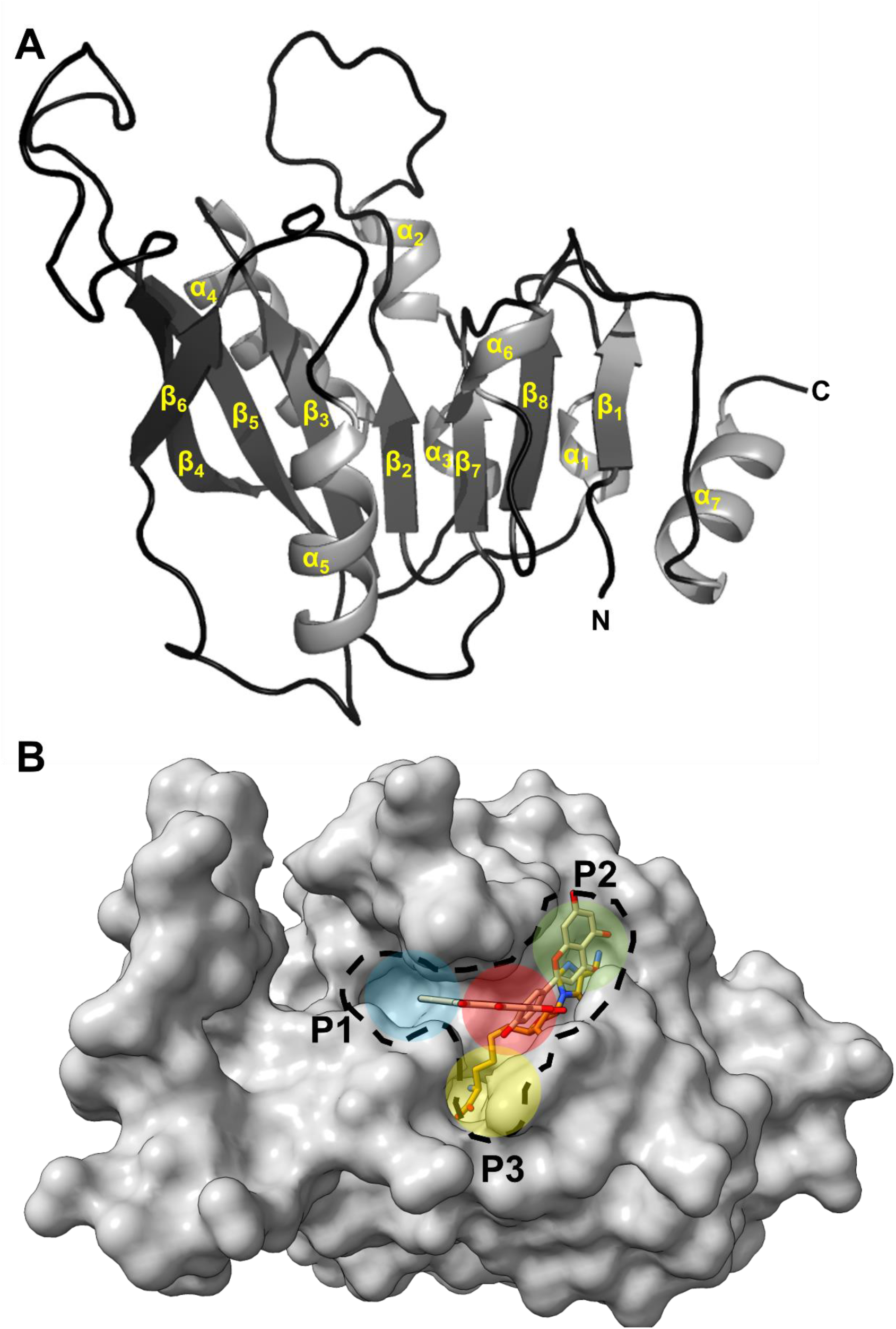
Crystal structure of METTL3 (PDB ID 5IL2), (A) The cartoon representation of human METTL3 is describing spatial arrangement of the secondary structure elements of the protein. The α helices and β strands are marked according to their position in amino acid sequence. (B) The surface representation of the protein in SAH (represented in sticks, colour: yellow) and AMF (represented in sticks, colour: grey) bound form depicts the how these two ligand molecules are differentially occupying the Y shaped binding pocket annotated as P1(blue), P2 (Green) and P3 (yellow). The centre of the Y shaped cavity is shown in red. SAH has occupied P2 & P3 while AMF binds to P1 & P2.

In this study an unbiased brute-force screening combined with de-novo drug designing approach directed by the binding pocket geometry has been applied to design highly specific inhibitors, that are druggable and easy to synthesize, against METTL3.

## RESULTS and DISCUSSION

### Molecular docking

In this study, a brute force approach was taken to screen over 3600 compounds (a library of commercially available compounds and FDA approved drugs with available 3D conformers) by molecular docking using Autodock Vina (ADV), to find suitable leads for rational drug design against METTL3. To validate the docking protocol, used in this screening, the receptor was docked with SAH, which had a docking score, expressed in kcal/mol, annotated here as ADV energy of -8.7kcal/mol (-36.4 kJ/mol). The RMSD of the docked pose of SAH compared to the SAH bound crystal structure of METTL3 (PDB ID: 5IL2) was 0.097nm (calculated using <u>DockRMSD: Docking Pose Distance Calculation (zhanggroup.org)</u> (Bell & Zhang, 2019). The results of this brute force screening were preliminarily sorted on the basis of ADV energy. The ADV energy of docked SAH (-36.4 kJ/mol) was considered as a cut off and nearly 250 drug molecules were found to have significantly greater binding. The ADV energy, from docking, of the top 10 compounds ranged from -12.1kcal/mol (-50.6kJ/mol) to -10.1kcal/mol (-42.25kJ/mol), which is significantly high compared to the ADV energy of SAH. The respective docking score of the top 10 compounds are tabulated in Figure 2 with their chemical structure and each compound was assigned a three-letter code for convenience, and these codes will be used for further references in the following text.

**Figure 2:**
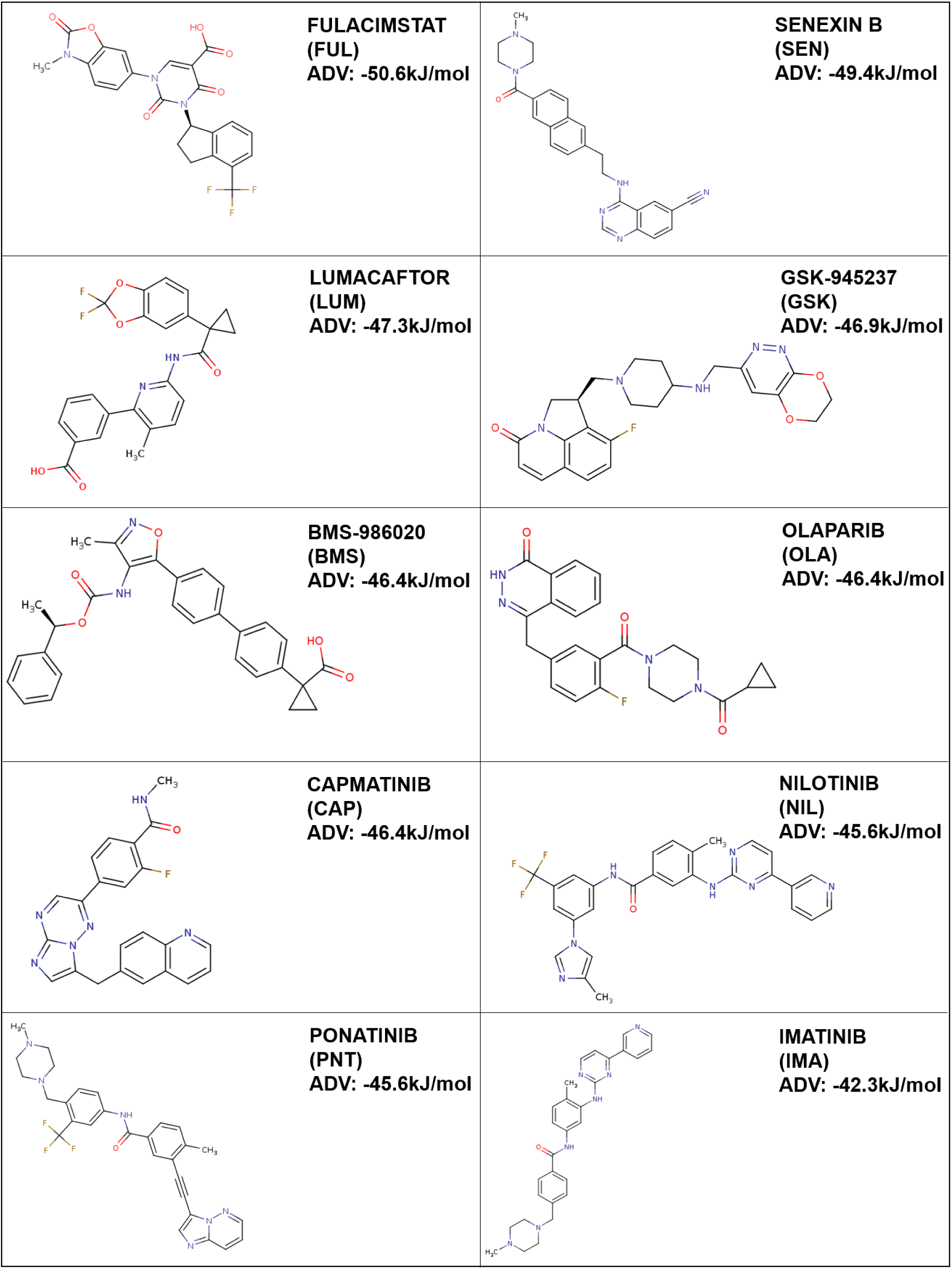
Chemical structure, name and the three-letter code of the top 10 molecules according to ADV energy.

### ADME and Toxicity Prediction

The druggability and toxicity of the top 10 compounds based on ADV energy was estimated using SWISS-ADME (Daina et al., 2017). Druggability of these compounds were assessed following Lipinski′s rule of five, i.e molecular weight, n-Octanol partition coefficient, number of hydrogen bond donors and acceptors. According to Lipinski′s rule, the top 10 compounds were druggable (Supplementary Table 1). The toxicity of the compounds, were also assessed using Pro Tox-II server (Banerjee et al., 2018). Which indicated that FUL, SEN, LUM, BMS, OLA, NIL belongs to class 4, whereas, GSK, CAP & IMA fall under class 3 and PNT fall under class 5 where higher the class, lower the toxicity, where class 1 is extremely toxic and class 6 being non-toxic.

### Molecular Dynamics Simulation

The top ten compounds based on ADV energy (Figure 2) were subjected to molecular dynamics simulation (MD simulation) followed by binding free energy calculation to perform second round of screening for finding the stronger interactions among the top 10 compounds. To achieve this goal, root mean square deviation (RMSD), root mean square fluctuation (RMSF), solvent accessible surface area (SASA) and radius of gyration (R_g_) were calculated from the MD simulation trajectories and the average values of RMSD, RMSF, SASA and Rg of the different ligand bound complexes are tabulated in Table 1.

**Table 1:** Average Root Mean Square Deviation (RMSD), Root Mean Square Fluctuation (RMSF), Radius of Gyration (Rg), Solvent Accessible Surface Area (SASA)

| Sr. No. | Compound Code | Average RMSD (nm) | Average RMSF (nm) | Average Rg (nm) | Average SASA (nm<sup>2</sup>) |
| --- | --- | --- | --- | --- | --- |
| 1 | FUL | 0.225 | 0.093 | 1.732 | 113.993 |
| 2 | SEN | 0.207 | 0.085 | 1.738 | 117.363 |
| 3 | LUM | 0.279 | 0.114 | 1.734 | 116.133 |
| 4 | GSK | 0.224 | 0.089 | 1.729 | 113.521 |
| 5 | BMS | 0.235 | 0.117 | 1.752 | 118.338 |
| 6 | OLA | 0.207 | 0.100 | 1.721 | 113.414 |
| 7 | CAP | 0.234 | 0.116 | 1.755 | 120.666 |
| 8 | NIL | 0.241 | 0.107 | 1.731 | 115.410 |
| 9 | PNT | 0.187 | 0.097 | 1.743 | 117.944 |
| 10 | IMA | 0.218 | 0.097 | 1.724 | 115.267 |

**RMSD:** The average RMSD of the apo-protein is 0.22nm. The overall average RMSD of the protein bound to different inhibitors is comparable to the average RMSD of the protein alone, ranging from 0.19nm to

0.24 nm, except LUM that has a slightly higher RMSD of 0.28nm. The change of RMSD value of the protein backbone for different inhibitor-protein complexes as well as for the apo-protein was compared over the course of simulation by plotting the RMSD against time in ns (Figure 3A).

**Figure 3:**
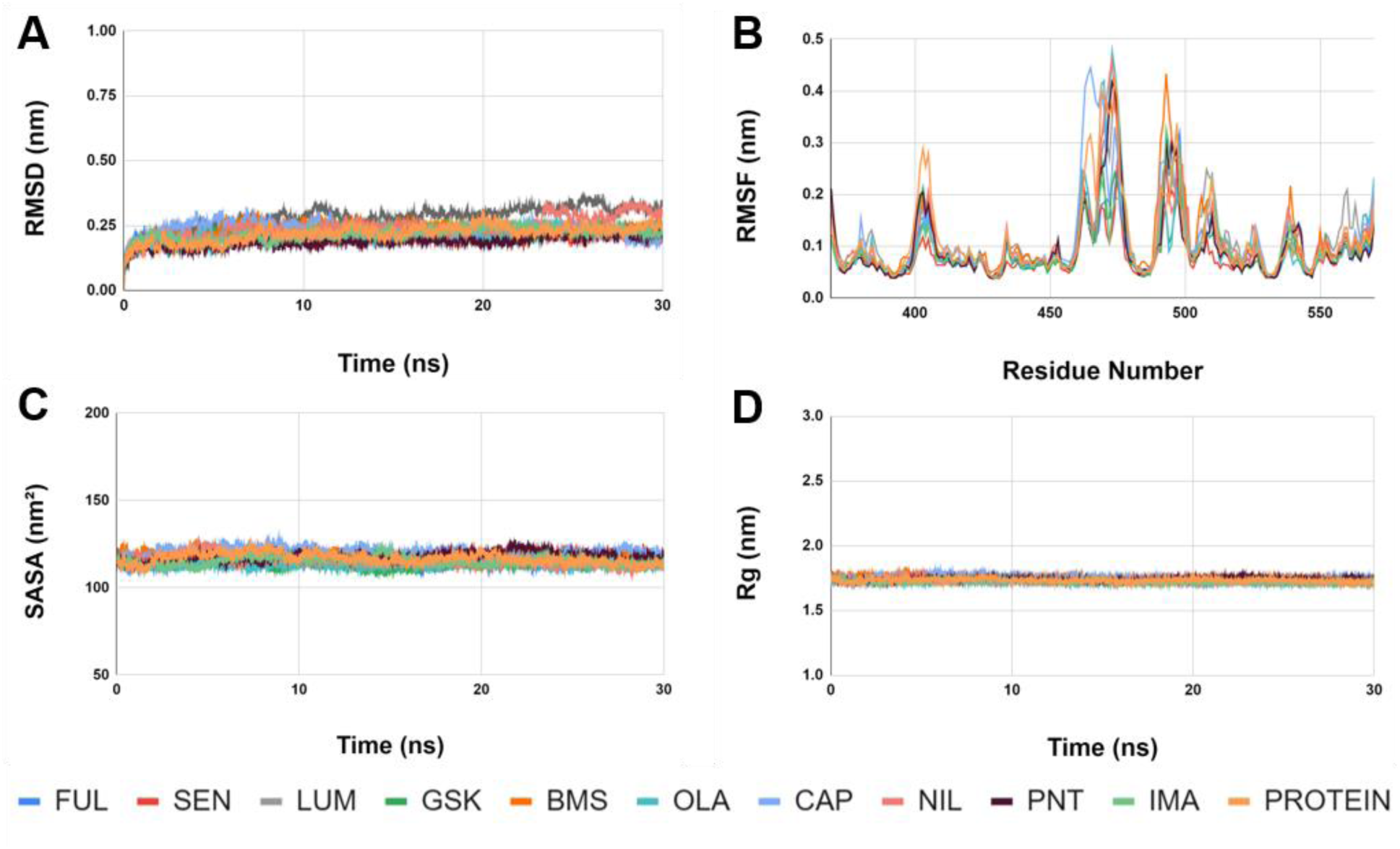
Representation of the molecular dynamics simulation results of the METTL3 alone as well as in the inhibitor bound form. The inhibitor bound protein complexes and the protein alone are shown in different colours. (A) The RMSD (nm) of backbone vs time in ns (B) backbone RMSF (nm) against the amino residue number, (C) solvent accessible surface area (SASA) in nm^2^ vs time (ns) and (D) Radius of gyration (R_g_) in nm vs time in ns.

**RMSF:** RMSF is the measure of flexibility of amino acid residues throughout the simulation process. The fluctuation of residues is calculated in the presence and absence of different ligand molecules. The RMSF of protein alone is 0.114nm, the RMSF of protein-inhibitor complexes range from 0.085nm - 0.117nm (Figure 3B). The higher fluctuation observed in RMSF plot around the residue 465 and 490 is due to the presence of two large loops residing between β_4_ and β_5_ (residue 460 – 480) and connecting β_5_ with β_6_ (residue 488 – 502) (Figure 1A).

**SASA**: The solvent accessible surface area (SASA) of the protein alone is 116.82nm ^2^. The SASA value for inhibitor bound complexes, varies from 113.52nm^2^ to 120.67nm^2^ (Figure 3C).

**Radius of gyration:** The compactness of the protein is indicated by radius of gyration (R_g_). The average R_g_ value for the protein is 1.73nm. For the average R_g_ ranges from 1.72-1.76nm, which is comparable with the protein alone, indicating no major structural changes happen during the course of simulation and the protein remains compact throughout the time course (Figure 3D).

### Protein ligand interaction

To determine whether the interactions between the inhibitor compounds and the protein throughout the course of simulation were governed by hydrogen bond or hydrophobic interaction, the co-ordinates of each inhibitor-protein complex were extracted every 5ns and analysed by Ligplot (Wallace et al., 1995). Figure 4 shows the result where the contribution from hydrogen bond and hydrophobic interaction has been expressed in terms of percentage. It is evident that hydrophobic interaction is the dominating force in the dynamics of protein-ligand interaction. The contribution of Lennard-Jones potential and Coulombic Short range interaction energy also portrays that the contribution from van der Waals interaction is greater than the electrostatic (Supplementary Figure 1).

**Figure 4:**
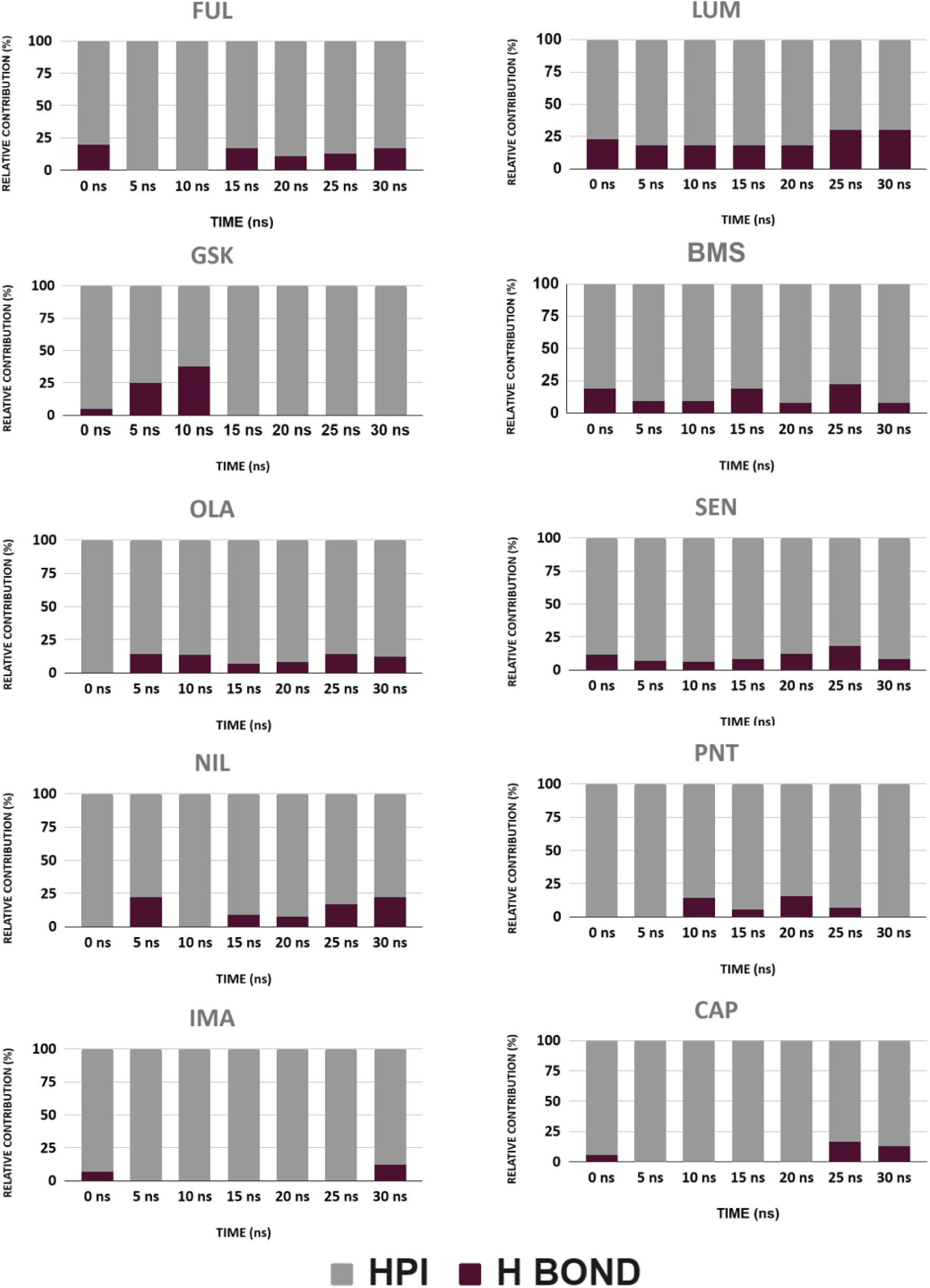
This stacked bar diagram represents the relative contribution of hydrophobic interaction (HPI) and hydrogen bond (H BOND) in protein-inhibitor interaction over the time period of MD simulation.

### MM-PBSA Calculation

The binding free energy (ΔG) of the protein-small molecule complexes of FUL, SEN, LUM, GSK, BMS, OLA, CAP, NIL, PNT & IMA, (top 10 compounds from docking) were calculated from the last 10ns trajectories of MD simulation. Among these, SEN and CAP failed to show significant interaction in terms of binding free energy, hence were rejected for further analysis. The binding poses of these remaining 8 compounds in the “Y” shaped binding pocket along with the interacting residues are presented in Figure 5. It is evident that all of the compounds have occupied sub-pocket P1 & P2 as described in the introduction and shows good agreement with the results of previous studies (Manna et al., 2023).

**Figure 5:**
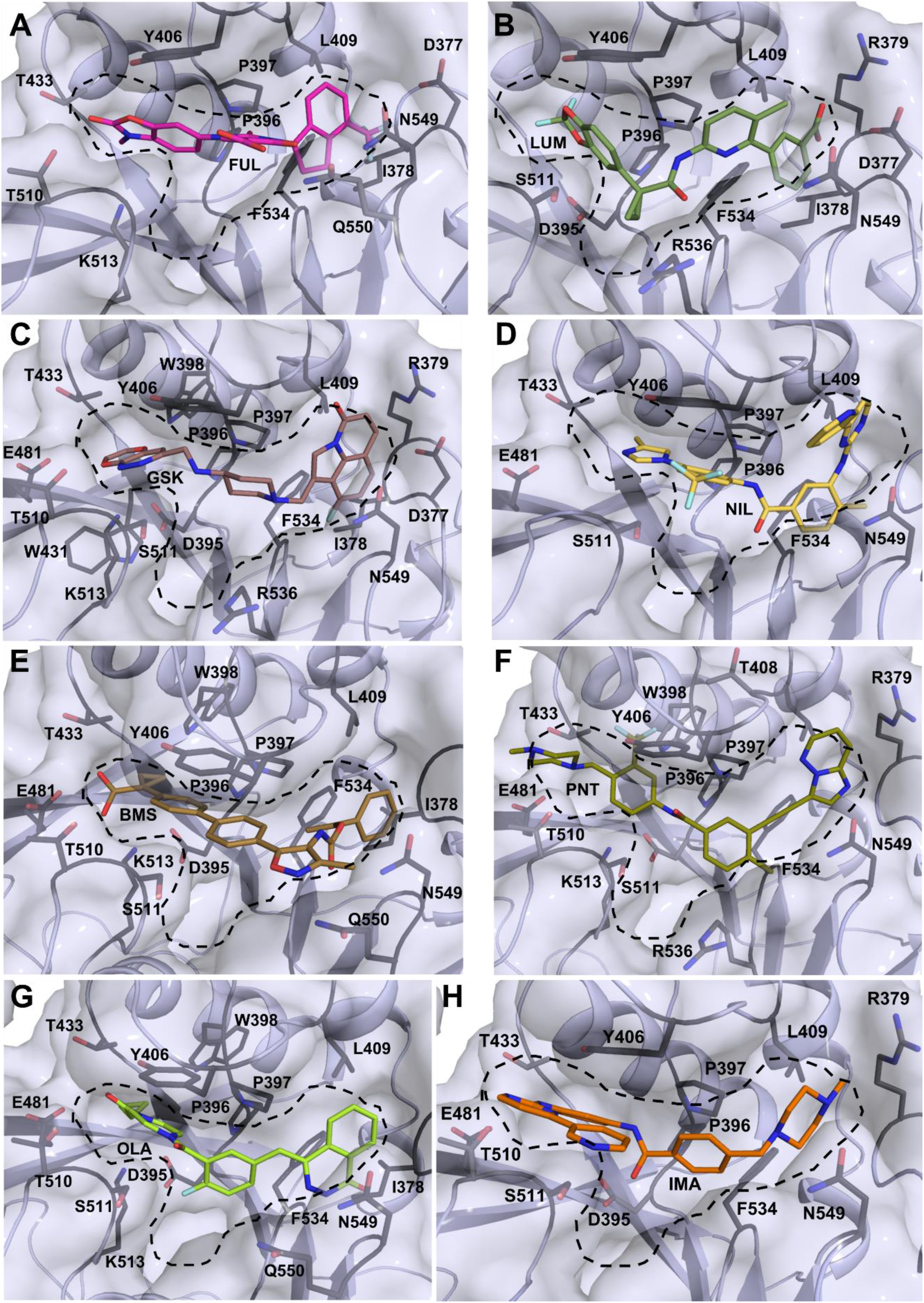
The residue level interaction between METTL3 and docked inhibitors shown in sticks, (A) FUL, (B) LUM, (C) GSK, (D) NIL, (E) BMS, (F) PNT, (G) OLA and (H) IMA revealing the binding pose of top 8 ligand molecules in the Y shaped binding pocket. The interacting residues of protein are shown in sticks and labelled with the three-letter code.

BMS, NIL, LUM, IMA and OLA have highly favourable binding (<-55kJ/mol), calculated by MM-PBSA method (Figure 6A). When the binding free energy contribution was further dissected into Van der Waals force of interaction and electrostatic force of interaction, portrayed in Figure 6B, it showed that Van der Waals force of interaction has a much higher contribution in binding of the inhibitor compounds in the P1-P2 cavity of the Y shaped pocket than the electrostatic force of interaction.

**Figure 6:**
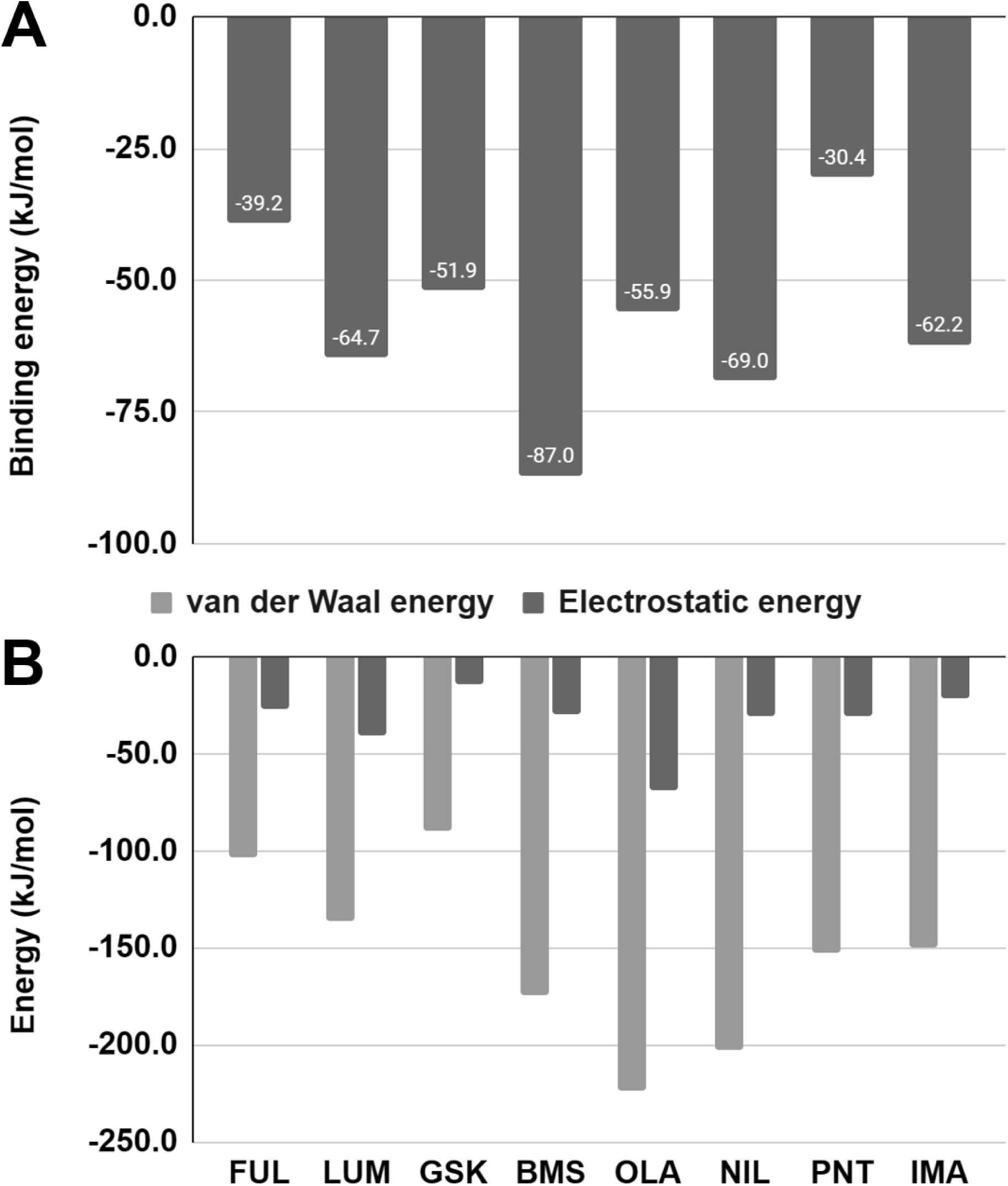
MM-PBSA based binding free energy. (A) ΔG values for METTL3-inhibitor binding expressed in kJ/mol. (B) The relative contribution of van der Waal’s energy and electrostatic energy in binding, also expressed in kJ/mol.

As discussed in the introduction section SAM/SAH binding pocket of METTL3 has 3 arms, P1, P2 and P3. SAM and SAH occupies P2 and P3. Whereas the docked compounds along with STM2457 occupy P1 and P2, leaving the P3 unoccupied. It inspired the authors in taking advantage of this unique Y shaped pocket to design Y shaped molecules that will fit in the SAM/SAH binding pocket (P2-P3) as well as P1 and become highly specific inhibitor of METTL3.

### Rational drug design

The binding poses of the top compounds, BMS, NIL, LUM, IMA, OLA as determined from docking, MD simulation and MM-PBSA calculation, along with AMF, RAD, MEH (reported as potent inhibitors of METTL3 in earlier study by the authors) (Manna et al., 2023)were assessed and used for structure guided de-novo drug designing to construct inhibitor molecules specific for the Y shaped binding pocket. The Y-shaped cavity of METTL3 protein can potentially host a branched, similarly Y-shaped molecule with 3 arms, called ′spokes′, connected at one end to a central ′hub′ group. The free termini of the spokes are meant to be so functionalized that they can effectively anchor that arm of the molecule in one of the sub-pockets (P1, P2 or P3) of METTL3 (Figure 1B). Thus, chemical functionalization of the spoke termini renders them selective of the pocket they occupy. The authors took help of structure guided de-novo drug designing approach to generate a series of molecules where successive cycles of new molecule designing and docking were performed to validate the course of rational drug designing, discussed in details in the following section. Careful analysis of SAM and SAH bound structures revealed that NH_2_-CHR-COOH group (where R refers to alkylic or phenylic groups) always occupies P3. Similarly, sub-pockets P1 and P2 show an affinity for OH group and CF_3_ group respectively (Figure 7). This is established in the binding poses of FUL, GSK, RAD where a CF_3_ group or C-F bonds are found interacting with P2 as well as in compounds like AMF where OH groups are found interacting with P1. However, the selectivity of OH for P1 and that of CF_3_ for P2 is not so strict, as the reverse cases where OH groups occupy P2 and CF_3_ occupies P1 are also noted (eg: AMF, RAD) (Figure 7) but never P3. P2 shows affinity for hydrophobic groups also, as illustrated by the binding modes of BMS, GSK and NIL. In most of the compounds, occupying P1& P2, like AMF, FUL, NIL, BMS, RAD, a central heterocyclic ring is widely conserved, addressed here as the central hub,

**Figure 7:**
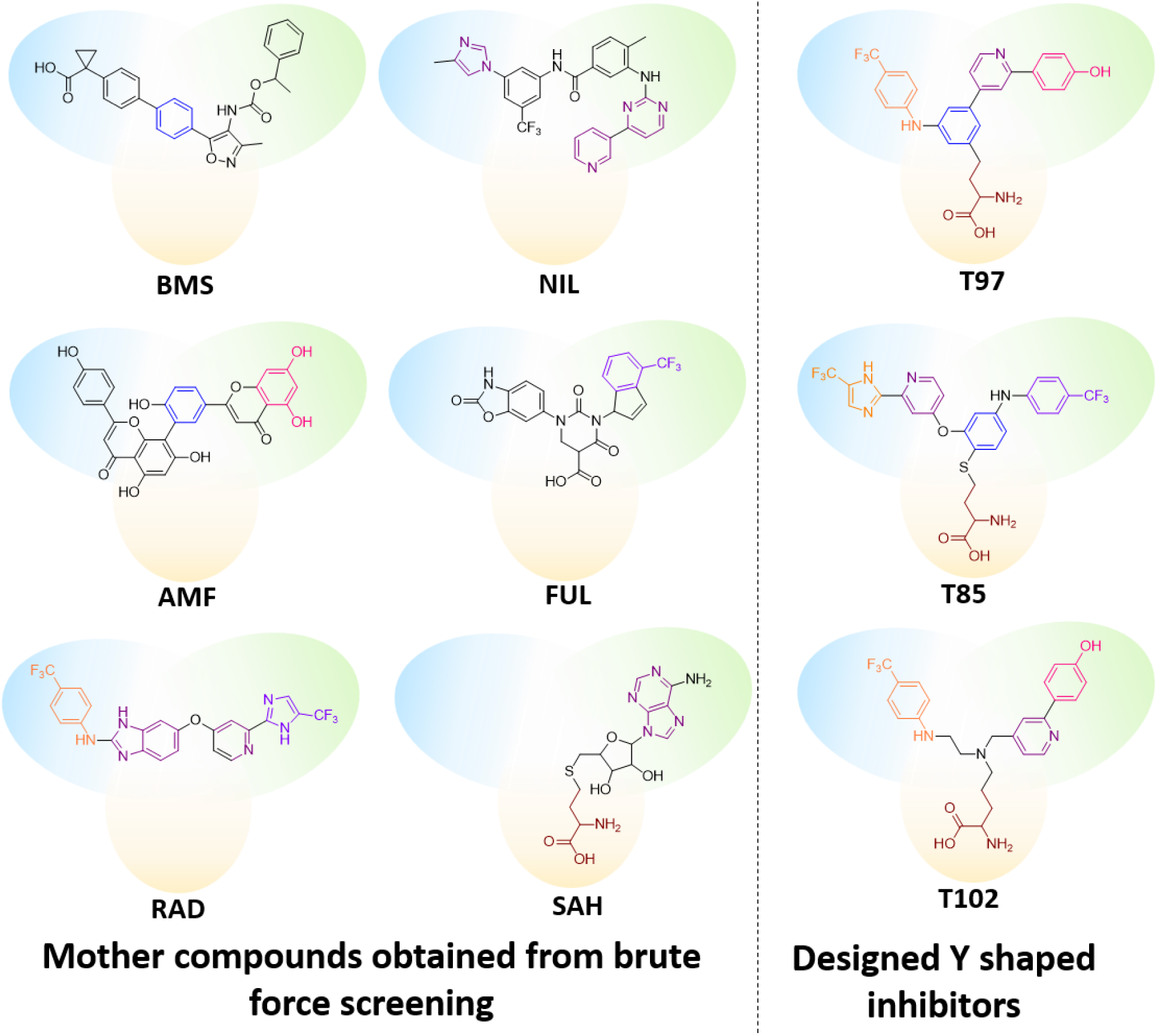
Graphical representation of de novo drug design strategy. Structural fragments derived from mother compounds (left) that are present in the designed inhibitors (right) are highlighted in the same colour (orange, pink, maroon, purple, magenta and royal blue). The protein sub-pockets P1, P2 and P3 are represented by blue, green and yellow halos respectively. The pictographic representation illustrates how the compounds fit in the METTL3 cavity.

anchors the molecule via strong stacking interaction with TYR406. For better understanding the binding pose of three ligands, AMF, NIL, SAH are visualized in Figure 8A, B, C respectively, where AMF and NIL occupies P1-P2 and SAH occupies P2-P3. To further bolster the notion that certain structural units or functional groups have a bias towards certain sub-pockets, when a glycine molecule (NH_2_-CH_2_-COOH), is docked into the cavity of METTL3, it occupies the binding pocket P3 (Figure 8D). Docking phenol (C_6_H_5_-OH), 1,3,5-trihydroxybenzene (C₆H₃(OH)₃) and trifluoro toluene (C_6_H_5_-CF_3_) (not shown in the figure) shows that either of them can dock in either P1 or P2 but not P3 (Figure 8E, 8F). Incidentally, docking a completely hydrophobic group, like benzene, in METTL3 shows that the molecule can occupy P2 as well as the intersectional region of the Y-shaped cavity. Thus, for the designed three-spoked molecules, one of the three termini always contains the NH_2_-CHR-COOH group to help the molecule bind to the P3 sub-pocket of the binding cavity. The other two termini are functionalised with a combination of phenolic OH groups, CF_3_ or hydrophobic phenyl groups as either can help the molecules to bind in P1 and P2 sub-pockets. The central hub is composed of a 1,3,5-trisubstituted phenyl, 1,2,5-trisubstituted phenyl, CHX_3_ groups (X = R, OR, NHR, -CH_2_-) or NR_3_. The feature setting the designed molecules apart from the compounds in brute-force trial is that they are all branched to fit the Y-shape of the binding cavity of METTLE3, like fingers in a glove. This combined with appropriate functionalisation improves the specificity and binding affinity of the compounds.

**Figure 8:**
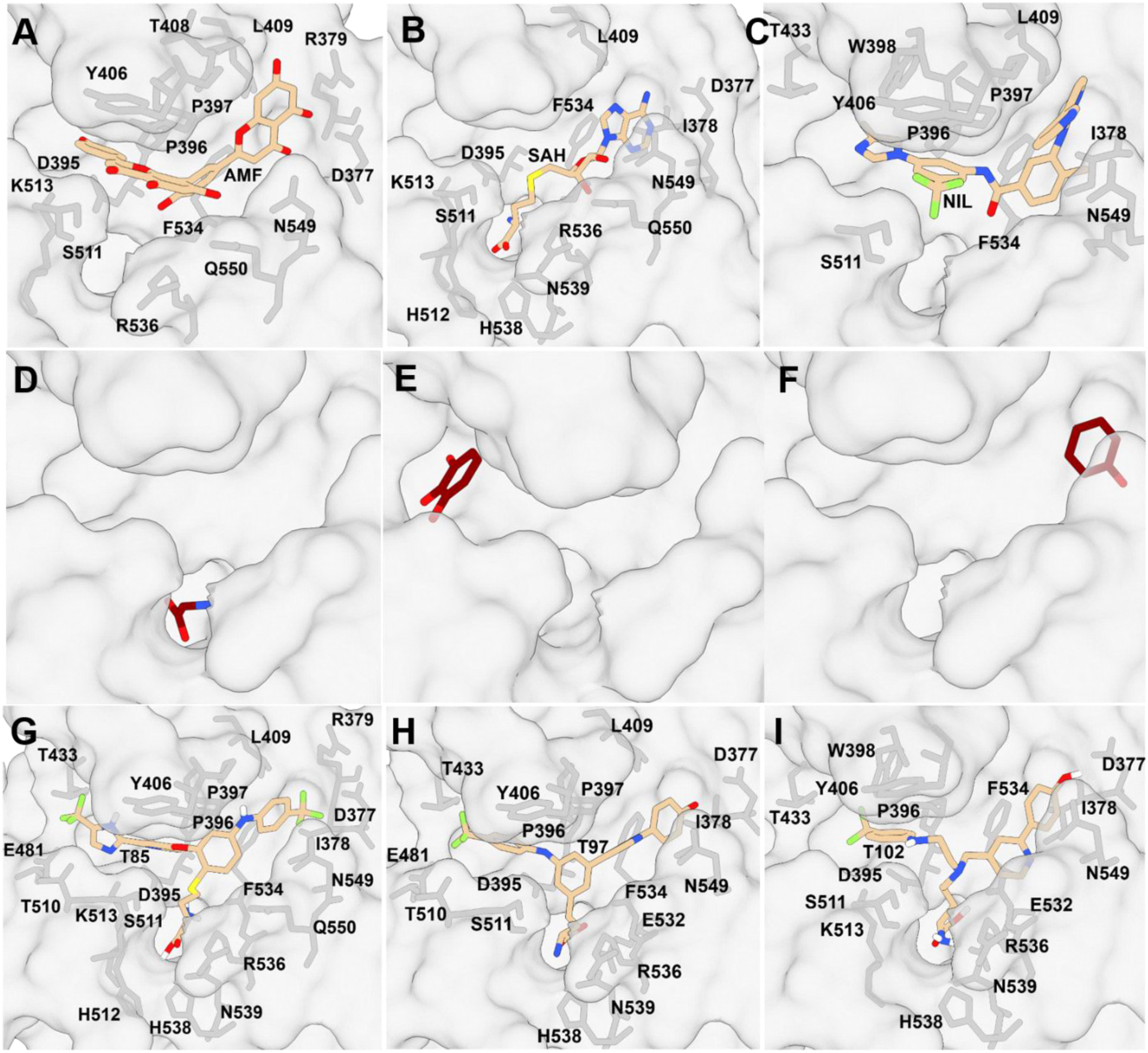
The upper panel shows binding pose of AMF (A) SAH (B) and NIL (C) in the Y shaped pocket where the ligands (inhibitor molecules) and interacting amino acid side chains and shown in stick whereas the receptor protein is presented in surface mode. Middle panel describes strategy of designing new molecules where three small chemical groups, Trihydroxybenzene (D), glycine (E) and phenol (F) have occupied three different pockets. The lower panel shows binding pose T85 (G), T97 (H) and T102 (I) in the Y shaped pocket along with the interacting amino acid side chains. T85, T97 and T102 occupies all of the three binding pockets (P1, P2 & P3) where they show additional interactions with amino acids H512 and H538 similar to SAH.

### Validation of the designed inhibitors

Using this strategy 125 Y shaped molecules were designed with variable spokes for trials (annotated as TR, where T stands for trial and R is the trial number expressed in alpha numeric code). These molecules were docked using ADV following the same protocol. The compounds with highest docking score were further screened based on the synthetic accessibility and druggability calculated from SwissADME. These Y shaped molecules are occupying the P1-P2-P3, as shown in Figure 8 G, H, I. The chemical structures of the top 10 (based on ADV energy) designed molecules are presented in Figure 9 along with ADV energy and synthetic accessibility score. 8 Y shaped molecules were selected based on their fit in the binding pocket and subjected MD simulation followed by binding free energy calculation using MM-PBSA method for the final round of selection. 3 compounds, T85, T97 & T102 have binding free energy better than -60kJ/mol (Supplementary Figure 2). The MD simulation data (Figure 10) clearly carries the signs of a stable protein-inhibitor complex during the run. The METTL3 protein with the top 2 compounds from the list, T85 and T97 were further simulated for 100ns to check whether the METTL3-inhibitor complex is stable in longer period of simulation. The output data shown in Figure 11 and Table 2 indicates the presence of a stable complex in terms of RMSD, RMSF, SASA and Rg. The binding free energy calculated from the last 10ns trajectories also reveals favourable binding and the binding free energy is in good agreement with the previous MD run of shorter run time.

**Figure 9:**
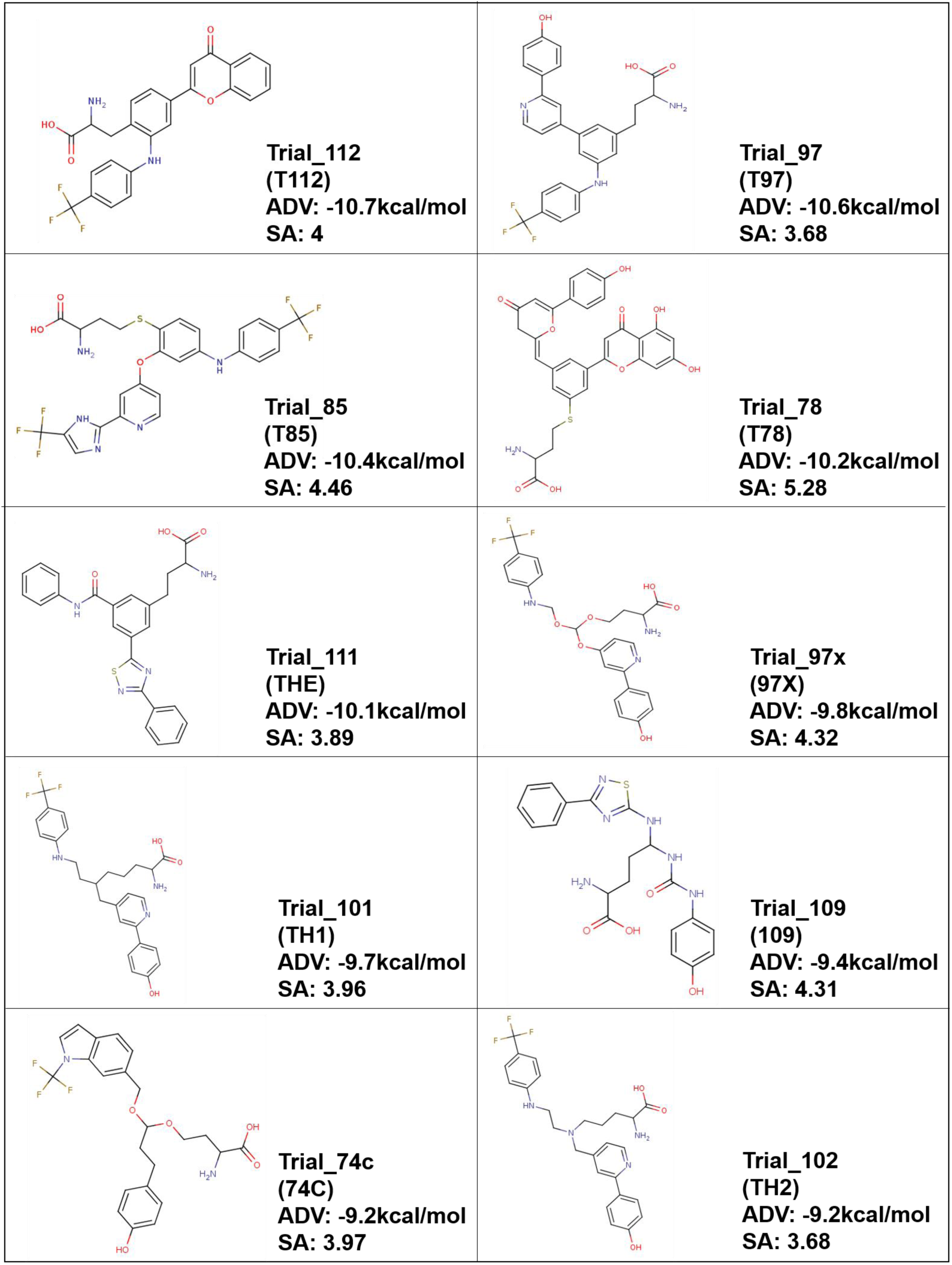
Chemical structure, the three-letter code, synthetic accessibility score of the top 10, Y shaped molecules designed for this study along with the trial number and ADV energy. For synthetic accessibility, 1: very easy to synthesize, 10: very difficult to synthesize.

**Figure 10:**
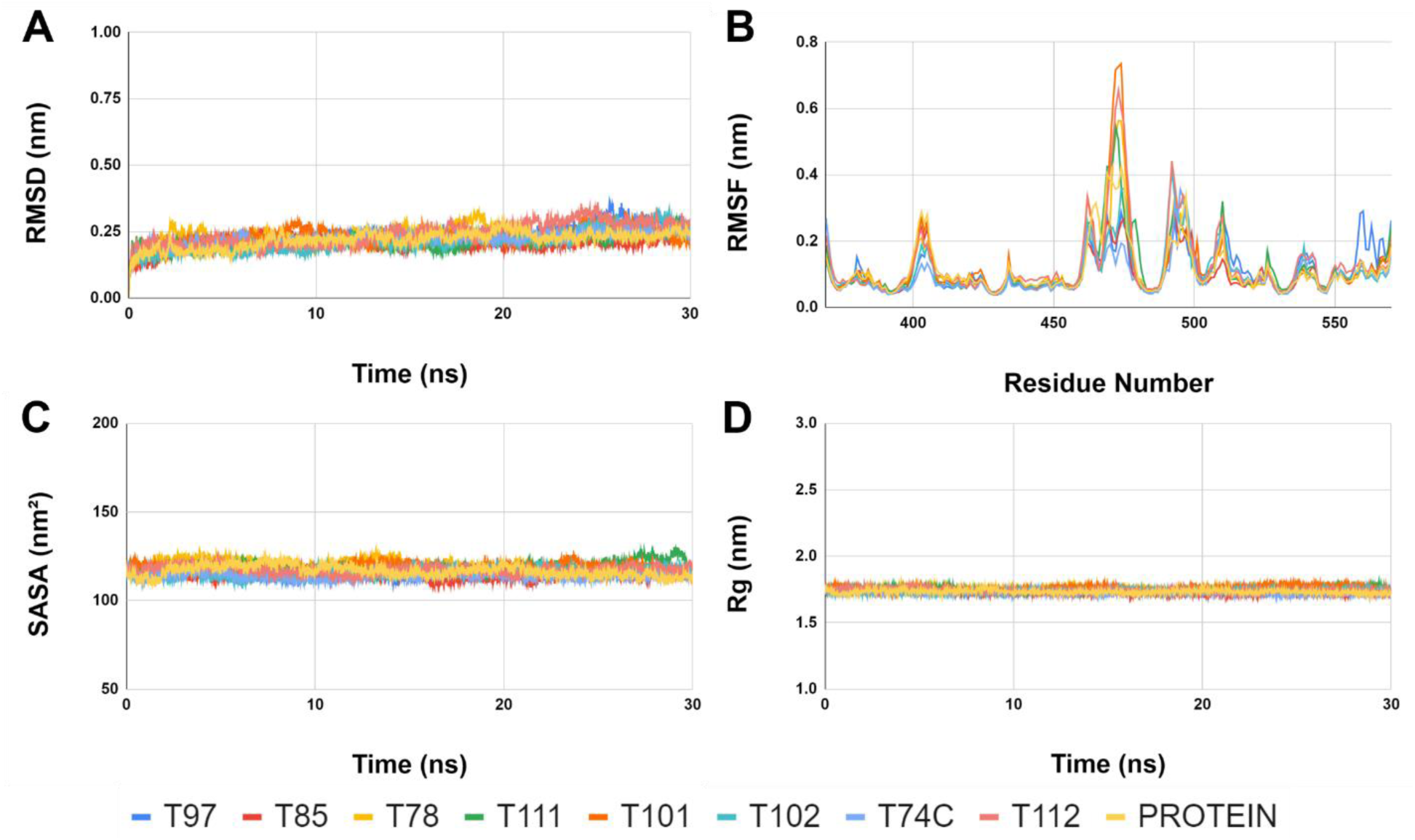
Representation of the molecular dynamics simulation results of the METTL3 alone as well as in the Y shaped inhibitor bound form. The inhibitor bound protein complexes and the protein alone are shown in different colours. (A) The RMSD (nm) of backbone vs time in ns (B) backbone RMSF (nm) against the amino residue number, (C) solvent accessible surface area (SASA) in nm^2^ vs time (ns) and (D) Radius of gyration (R_g_) in nm vs time in ns.

**Figure 11:**
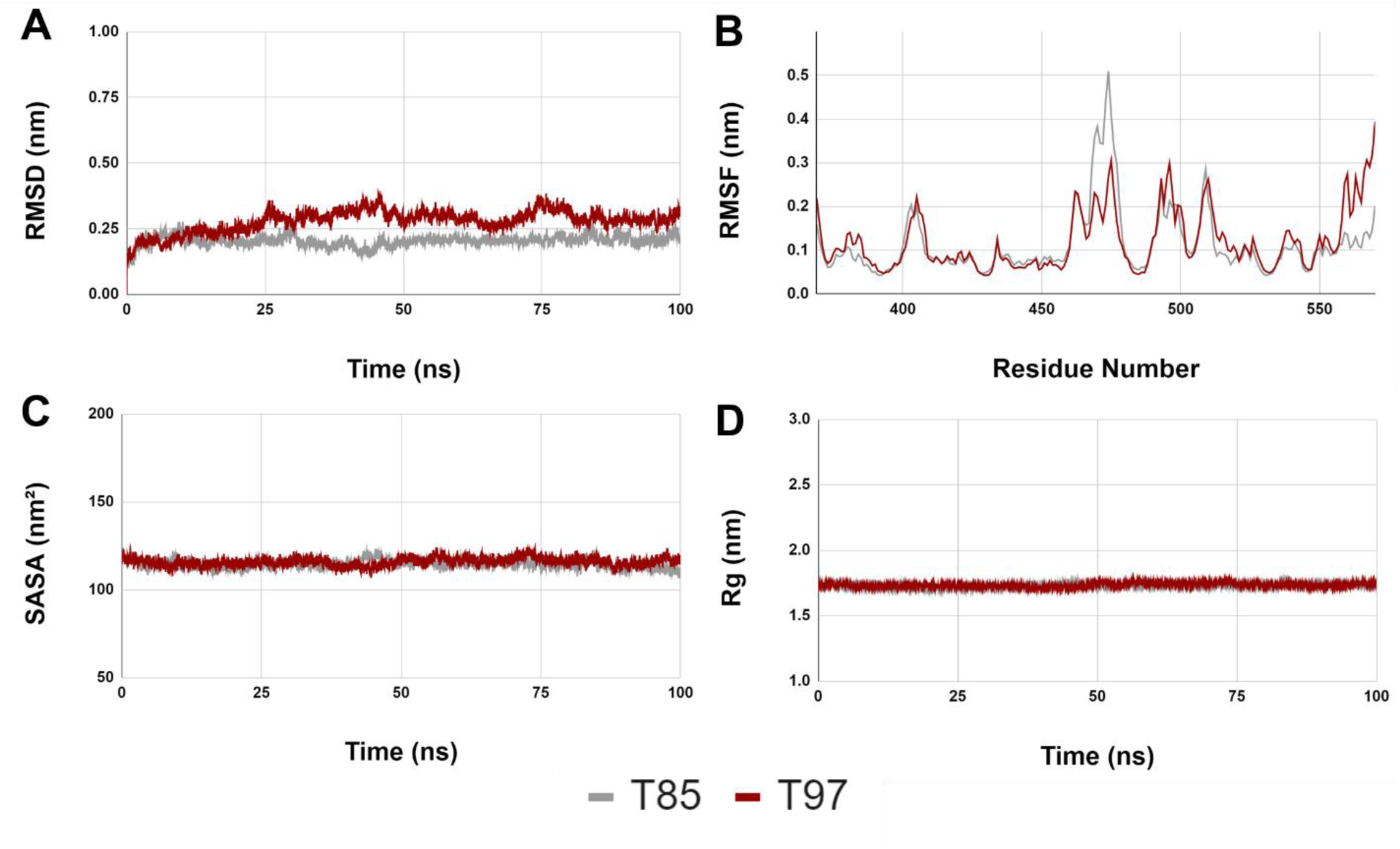
Representation the stability of protein-inhibitor complex (T97 and T85) during 100ns MD simulation. The MD simulation output is presented in (A) The RMSD (nm) of backbone vs time in ns, (B) backbone RMSF (nm) against the amino residue number, (C) solvent accessible surface area (SASA) in nm^2^ vs time (ns) and (D) Radius of gyration (R_g_) in nm vs time in ns.

**Table 2:**
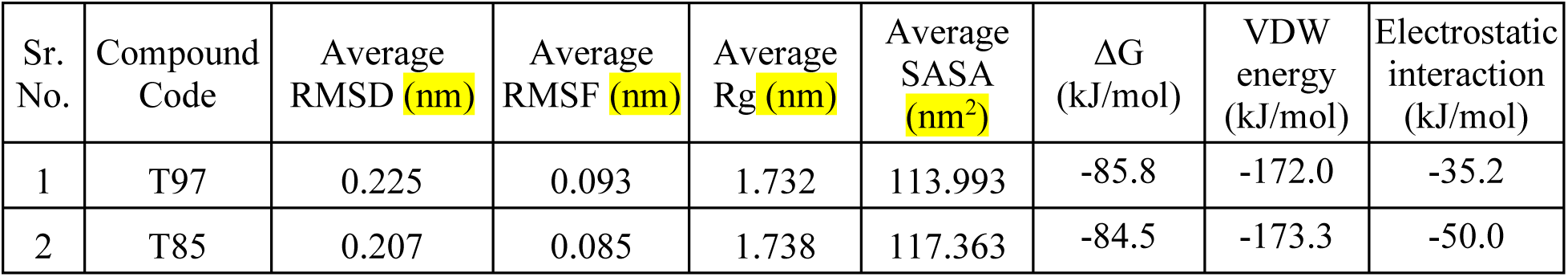
Average Root Mean Square Deviation (RMSD), Root Mean Square Fluctuation (RMSF), Radius of Gyration (Rg), Solvent Accessible Surface Area (SASA), protein ligand binding free energy (ΔG), van der Waal’s energy and electrostatic energy.

## Conclusion

Due to the unique structure these Y shaped molecules are highly specific inhibitors for METTL3. To assess this hypothesis, the Ki of the only commercially available inhibitor of METTL3, STM2457, was compared to the Ki of T97, T85, T102, determined by Autodock 4.2 (Morris et al., 2009). It has been found that the Ki of T97 (5.74pM) and T85 (337.44pM) are comparable to the Ki of STM2457 (106.54pM) which is a significant improvement in terms of Ki, if we compare it with the Ki of the top compounds found from docking in this study as well as the previous study: NIL (18.36nM), AMF (596.66nM), MEH (3.02µM).

The highest binding free energy is shown by T97 in which the terminal functionalised by CF_3_ interacts with P1 and that functionalised with the phenolic OH interacts with P2. By modifying the hub group of T97 from a 1,3,5-trisubstituted phenyl to NR_3_, T102 was obtained, which also shows high binding free energy in MD experiments. In T85, CF3 group is used in both termini to bind to P1 and P2. It should be noted that the groups that connect the hubs to the functionalised terminals have mostly been derived from the known selection of compounds as well. In some cases, however, these groups have undergone extensive modification through trial and error to ensure better fit in the binding site as well as maximising interaction in the pockets.

This entire study was focused on designing highly specific inhibitor of METTL3 that could have the potency to be a good drug as be useful in epitranscriptomics research where catalytically silent METTL3 is extremely important to study its function. A specific inhibitor can serve this purpose well and make the experiments easier. Here authors took a unique approach by combining brute force screening and structure guided drug designing validated by iterative cycles of de novo drug designing, molecular docking and MD simulation based binding free energy calculation. Through this rigorous method of elimination after screening 3600 compound authors selected 8 compounds on the basis of docking score and binding free energy which were used as skeleton to designed 125 Y shaped compounds. From this pool 8 Y shaped compounds were further selected based on docking score, druggability, synthetic accessibility and finally subjected to MD simulation followed by MM-PBSA based energy calculation. In this round of screening, 3 best compounds were selected from MD simulation and MM-PBSA energy calculation. In the final round, based on Ki, authors are reporting two best Y shaped inhibitors, T97 & T85 amongst many more where structure guided de novo drug designing have constructed druggable, easy to synthesize compounds as potential drugs for AML, NSLC, etc.

## Materials and Methods

### Molecular docking and Brute force screening

For brute force screening Autodock Vina from PyRx platform (Dallakyan & Olson, 2015) was used as the work engine to perform high throughput molecular docking. X-ray crystal structure of the target human protein, METTL3 (PDB ID: 5IL2) was retrieved from the Protein Data Bank (https://www.rcsb.org/) and the SAM/SAH binding pocket of METTL3 was targeted for molecular docking. The grid box was set by including the SAM/SAH binding residues, which is comprised of amino acids ASP377, ASP395, ASN539, GLU532, ARG536, HIS538, ASN549, & GLN550 at the centre with the following grid box dimension 22 × 21 × 21. A library of more than 3600 commercially available drug molecules, with available 3D coordinates in PDB file format was curated by the authors using several small molecule databases (eg: PUBCHEM, Zinc, EMBL), for the preliminary brute force screening.

The detailed protocol of receptor and ligand preparation which includes removal of lone pairs, solvents, crystallization artifacts and addition of missing hydrogen, and charge has described in previous work (Mitra et al., 2023).

### Drug-like properties and Toxicity prediction

The drug like properties of top 10 compounds based on ADV energy, were analysed using SwissADME server (Daina et al., 2017). The toxicity was predicted by Pro Tox-II server.

### Molecular Dynamics simulation and Molecular Mechanics Poisson-Boltzmann Surface Area calculation protocol

The preliminarily selected drug molecules based on the ADV energy (Figure 2) in the protein-ligand complex form were subjected to Molecular dynamics simulation (MD Simulation) followed by Molecular Mechanics Poisson-Boltzmann Surface Area calculation (MMPBSA). All MD simulations were performed on Gromacs 5.1.2 (https://www.gromacs.org/) platform using CHARMM 36 force field (Vanommeslaeghe et al., 2009) and protein and ligand complex, was solvated using TIP3P water model (Jorgensen et al., 1983) in a dodecahedron box was used to perform. The ligand topology was generated using CGenFF server (https://cgenff.umaryland.edu/). The topology file of the solvated and electroneutral complex was subjected to energy minimization using the steepest decent algorithm keeping the force less than 10.0kJ/mol. Before final MD run the system was equilibrated by two stage equilibration process, NVT and NPT. NVT also known as isothermal-isochoric ensemble. NVT stands for number of particles (N), volume (V) and temperature (T). In this stage, the particles were kept constant at 300 K and maintained for 100 ps following the Berendsen′s method (Berendsen et al., 1984). The second equilibration phase is NPT or isothermal-isobaric ensemble, which has a constant pressure of 1 bar with an equilibration of number of particles (N), pressure (P) and temperature (T) for 100 ps following -Rahman barostat method (Parrinello & Rahman, 1981). Finally, the equilibrated system was subjected to MD run. The trajectories from MD simulation were corrected of periodic boundary conditions, and centred. From the corrected trajectories RMSD (Schreiner et al., 2012), RMSF (Maiorov & Crippen, 1994), R_g_ (Lobanov et al., 2008) and SASA (Lemkul, 2019) was calculated.

The last 10ns of corrected trajectory of each MD simulation experiment was extracted to calculate the protein-ligand binding free energy (ΔG_binding_) using MM-PBSA method with g-mmpbsa script g_mmpbsa (rashmikumari.github.io) (Kumari et al., 2014). The binding free energy was calculated using the following equation:

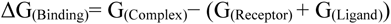

Where G_(Complex)_ denotes free energy of the protein-ligand complex, G_(Receptor)_ is the free energy of the receptor and G_(Ligand)_ is the free energy of the ligand in solvent.

The set of novel compounds, rationally designed in this study were also subjected to MD simulation and MMPBSA based binding free energy calculation using the protocol as described above.

### Rational designing of drug molecules: or Structure based drug designing

De novo ligand design involves utilizing computational tools to study the interaction of molecules with proper functional groups with a target protein. The molecules were designed using Chemical Sketch Tool from RCSB (<u>RCSB PDB: Chemical Sketch Tool</u>) and converted using OpenBabel (<u>OpenBabel</u>).

The geometries of the 5 designed molecules have additionally been optimized using Gaussian16c program package. Optimization and frequency calculation is carried out at B3LYP/TZVP(Becke, 1988, 1993; Lee et al., 1988) level for all atoms and the aqueous solvent environment is simulated using the polarization continuum model. The optimized geometries are characterized as minima on the potential energy surface of the molecules from the absence of any imaginary frequencies.

## Acknowledgement

The authors thank Ms Pragati Samal for helping in preliminary data acquisition.

## Statements & Declarations

### Contributions

All authors contributed to the study. All authors read and approved the final manuscript. The credit author statement is as follows:

DH and AR conceptualized the project, design the experiments, analyse the data, prepared the figures and wrote the manuscript. MG performed docking and MD simulation, tabulated the results, performed preliminary analysis and helped in manuscript writing. RG designed novel compounds, performed analysis and wrote the manuscript.

