## supplemental tables and figures for "De novo drug designing coupled with brute force screening and structure guided lead optimization gives highly specific inhibitor of METTL3: a potential cure for Acute Myeloid Leukaemia"

[a] Independent Researcher

[b] Department of Inorganic and Physical Chemistry, Indian Institute of Science, Bangalore, India

[c] Center for Healthcare Science and Technology, Indian Institute of Engineering Science and Technology, Shibpur, India

[d] Department of Biotechnology, St. Xavier's College, Kolkata, India

§ These authors contributed equally: Manisha Ganguly and Radhika Gupta

* Correspondence

**Supplementary Figure 1:** Coulombic interaction energy (Coul-SR) and the short-range Lennard-Jones energy (LJ-SR) expressed in kJ/mol

**Supplementary Figure 2:** (A) ΔG of inhibitor binding is expressed in kJ/mol, (B) van der Waal's energy and electrostatic energy for ligand binding expressed in kJ/mol.

**Supplementary Table 1:** SwissADME and Pro Tox-II analysis of top 10 compounds

Supplementary Figure 1

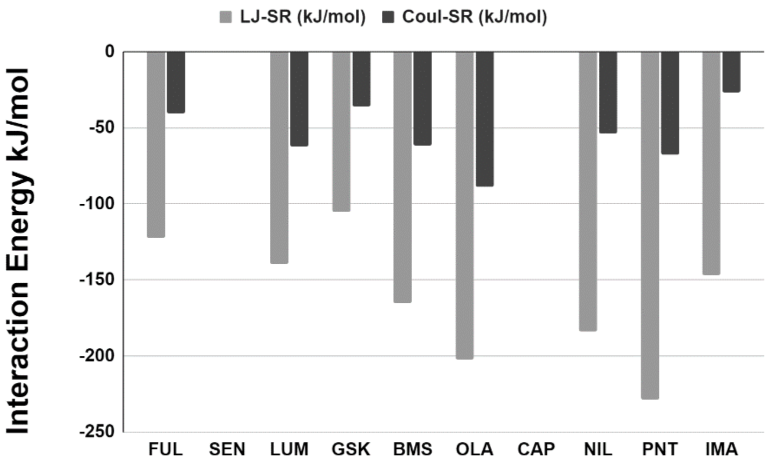


Supplementary Figure 2A

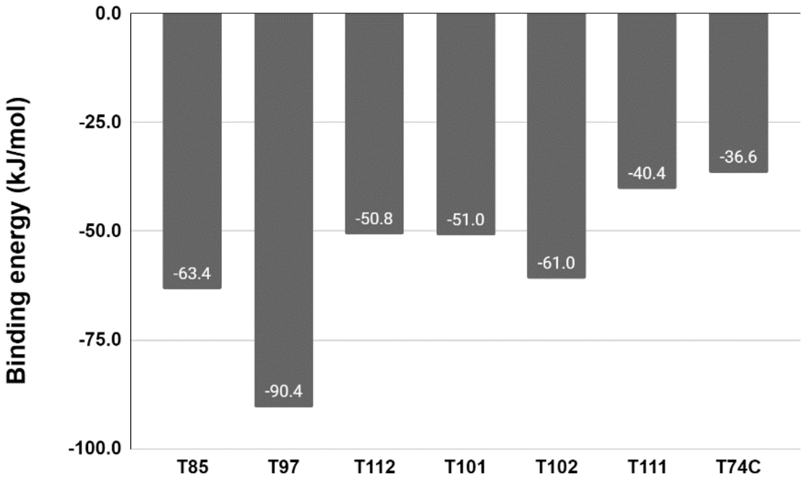


Supplementary Figure 2B

| Parameters | FUL | SEN | LUM | GSK | BMS | OLA | CAP | NIL | PNT | IMA |
|---|---|---|---|---|---|---|---|---|---|---|
| **MW** | 487.38 | 450.53 | 452.41 | 451.49 | 482.53 | 434.46 | 412.42 | 529.52 | 532.56 | 493.6 |
| **Acceptable range** | ≤ 500 | ≤ 500 | ≤ 500 | ≤ 500 | ≤ 500 | ≤ 500 | ≤ 500 | ≤ 500 | ≤ 500 | ≤ 500 |
| **NHBA** | 9 | 5 | 8 | 8 | 6 | 5 | 6 | 8 | 8 | 6 |
| **Acceptable range** | ≤ 10 | ≤ 10 | ≤ 10 | ≤ 10 | ≤ 10 | ≤ 10 | ≤ 10 | ≤ 10 | ≤ 10 | ≤ 10 |
| **NHBD** | 1 | 1 | 2 | 1 | 2 | 1 | 1 | 2 | 1 | 2 |
| **Acceptable range** | ≤ 5 | ≤ 5 | ≤ 5 | ≤ 5 | ≤ 5 | ≤ 5 | ≤ 5 | ≤ 5 | ≤ 5 | ≤ 5 |
| **MR** | 116.96 | 141.29 | 113.98 | 124.37 | 136.35 | 125.21 | 113.93 | 141.08 | 150.1 | 154.5 |
| **Acceptable range** | 40–130 | 40–130 | 40–130 | 40–130 | 40–130 | 40–130 | 40–130 | 40–130 | 40–130 | 40–130 |
| **Mlogp** | 3.17 | 2.54 | 3.09 | 2.31 | 3.72 | 3.09 | 2.8 | 2.75 | 3.9 | 2.15 |
| **Acceptable range** | ≤ 4.15 | ≤ 4.15 | ≤ 4.15 | ≤ 4.15 | ≤ 4.15 | ≤ 4.15 | ≤ 4.15 | ≤ 4.15 | ≤ 4.15 | ≤ 4.15 |
| **Druglikeness** | Yes; 0 Violation | Yes; 0 Violation | Yes; 0 Violation | Yes; 0 Violation | Yes; 0 Violation | Yes; 0 Violation | Yes; 0 Violation | Yes; 1 Violation | Yes; 1 Violation | Yes; 0 Violation |
| **ProTox class** | 4 | 4 | 4 | 3 | 4 | 4 | 3 | 4 | 5 | 3 |

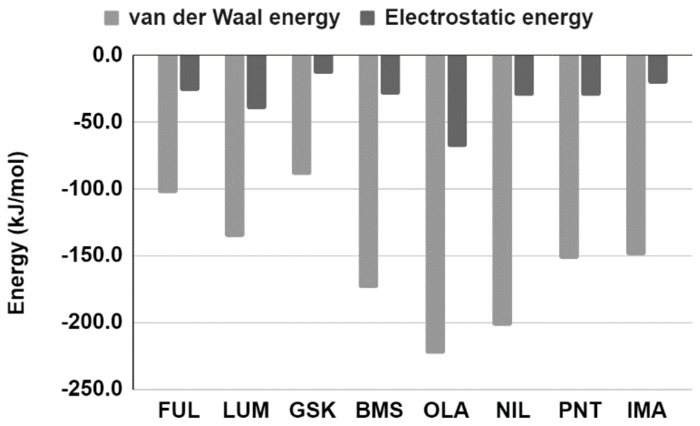


**Supplementary Table 1**
